# SARS-CoV-2 Spike protein promotes hyper-inflammatory response that can be ameliorated by Spike-antagonistic peptide and FDA-approved ER stress and MAP kinase inhibitors *in vitro*

**DOI:** 10.1101/2020.09.30.317818

**Authors:** Alan C-Y. Hsu, Guoqiang Wang, Andrew T. Reid, Punnam Chander Veerati, Prabuddha S. Pathinayake, Katie Daly, Jemma R. Mayall, Philip M. Hansbro, Jay C. Horvat, Fang Wang, Peter A. Wark

## Abstract

SARS-CoV-2 infection causes an inflammatory cytokine storm and acute lung injury. Currently there are no effective antiviral and/or anti-inflammatory therapies. Here we demonstrate that 2019 SARS-CoV-2 spike protein subunit 1 (CoV2-S1) induces high levels of NF-κB activations, production of pro-inflammatory cytokines and mild epithelial damage, in human bronchial epithelial cells. CoV2-S1-induced NF-κB activation requires S1 interaction with human ACE2 receptor and early activation of endoplasmic reticulum (ER) stress, and associated unfolded protein response (UPR), and MAP kinase signalling pathways. We developed an antagonistic peptide that inhibits S1-ACE2 interaction and CoV2-S1-induced productions of pro-inflammatory cytokines. The existing FDA-approved ER stress inhibitor, 4-phenylburic acid (4-PBA), and MAP kinase inhibitors, trametinib and ulixertinib, ameliorated CoV2-S1-induced inflammation and epithelial damage. These novel data highlight the potentials of peptide-based antivirals for novel ACE2-utilising CoVs, while repurposing existing drugs may be used as treatments to dampen elevated inflammation and lung injury mediated by SARS-CoV-2.

## Introduction

The emergence of a novel SARS-coronavirus in late 2019 (SARS-CoV-2; previously known as 2019-nCoV), and the CoV disease (COVID)-19 it causes, has led to a devastating pandemic of the 21^st^ century. SARS-CoV-2 belongs to the beta-coronavirus genus with approximately 79.5% sequence homology to the SARS-CoV that emerged in 2002 (Wang et al., 2020b). Similar to SARS-CoV (2002), this novel CoV also utilises angiotensin converting enzyme (ACE)2 as its host receptor to mediate membrane fusion and virus entry and viral replication (Zhou et al., 2020). SARS-CoV-2 spike protein contains two subunits, subunit 1 (S1) and S2, that mediates viral attachment to ACE2 and membrane fusion, respectively. The receptor-binding domain (RBD) of S1 is the critical region of the spike protein for ACE2 binding.

Human bronchial epithelial cells are both susceptible and permissive to CoV infection and replication, and the innate immune responses produced by which are critical in the early containment of infection and spread. Viral infection results in the activations of several pattern recognition receptors (PRRs) including retinoic acid-inducible gene I like receptors (RLRs; RIG-I and melanoma differentiation associated protein 5 (MDA-5))(Hayman et al., 2019; Saito and Gale, 2008), and toll-like receptors (TLRs; TLR3 and TLR7)(Alexopoulou et al., 2001; Diebold et al., 2004; Le Goffic et al., 2007). Recognition of viral RNAs by these PRRs leads to activation of transcription factor nuclear factor kappa-light-chain-enhancer of activated B cells (NF-κB), which facilitates the expression of pro-inflammatory cytokines such as interleukin (IL)-6 and IL-1β. These cytokines recruit and activate important immune cells including macrophages and neutrophils, which further promote inflammation and contain viral spread (Guan et al., 2020) (Wang et al., 2008). Patients with severe COVID-19 have been shown to have develop enhanced systemic inflammatory responses (aka. cytokine storm), and acute lung injury and acute respiratory distress syndrome (ARDS) (Huang et al., 2020; McGonagle et al., 2020; Mehta et al., 2020). This storm is characterised by heightened levels of IL-6, tumor necrosis factor-α (TNF-α), and C-C motif chemokine ligand (CCL)2. Currently there are no specific antiviral drugs available or anti-inflammatory drugs that have been shown to influence clinical outcomes for people with COVID-19. A broad-spectrum antiviral drug remdesivir is currently being used for COVID-19 through compassionate use requests as well as in clinical trials in the US and China. The anti-malarial drug hydroxychloroquine is also under several clinical trials, although preliminary reports have not shown beneficial effects (Magagnoli et al., 2020; Mahevas et al., 2020).

Here we demonstrate that the SARS-CoV-2 spike protein S1 (CoV2-S1) or RBD (CoV2-RBD) alone stimulates more pronounced production of pro-inflammatory cytokines as well as factors associated with epithelial damage, compared with the S1 and RBD of SARS-CoV (CoV-S1 and -RBD, respectively). This heightened inflammatory response requires S1 and ACE2 interaction and is primarily driven by early endoplasmic reticulum (ER) stress and its adaptive unfolded protein response (UPR), as well as activation of mitogen-activated protein (MAP) kinase signalling pathways. The early induction of ER-UPR results in activation of MAP kinase, and both pathways synergistically leads to NF-κB activation and production of the pro-inflammatory cytokines IL-6, IL-1β, TNF□, and CCL2, the latter three of which have all been shown to contribute to acute lung injury(Kolb et al., 2001) (Sheridan et al., 1997).

Since CoV2-S1 induces NF-κB activation *via* its interaction with ACE2 and early activations of ER-UPR and MAP kinase signallings, we designed a series of CoV2-S1-antagonistic peptides and identified a peptide, designated AP-6, that inhibits S1-ACE2 interaction. We show that CoV2-S1-mediated inflammatory response are not only ameliorated by AP-6, but the heightened inflammatory responses are also inhibited by an FDA-approved ER stress inhibitor 4-phenylbuic acid (4-PBA), and MAP kinase inhibitors trametinib (GSK1120212) and ulixertinib (BVD-523).

Taken together, these data shed light on how the spike protein of SARS-CoV-2 may contribute to the exaggerated inflammatory responses and pathology observed in those with severe COVID-19. The specific antagonistic peptide that specifically target S1 may serve as an antiviral against SARS-CoV-2. We also showed how existing, pathway specific, FDA-approved drugs could be re-purposed and immediately deployed to reduce infection-induced symptoms and pathologies that are primarily driven by exaggerated inflammatory responses.

## Results

### SARS-CoV-2 spike protein S1 and RBD induces elevated induction of pro-inflammatory responses

To investigate if SARS-CoV-2 spike protein induces production of pro-inflammatory cytokines, we used a minimally immortalised human bronchial epithelial cell line BCi-NS1.1, which was derived from human primary bronchial epithelial cells (Hayman et al., 2019; Hu et al., 2019; Kedzierski et al., 2017; Walters et al., 2013). BCi-NS1.1 was stimulated with histidine (His)-tagged CoV2-S1, CoV2-RBD, or CoV-S1 or CoV-RBD. Both S1 and RBD of CoV2 induced robust and similar levels of pro-inflammatory cytokines (IL-6, IL-1β, and TNF⍰) in a dose-dependent manner at 24 hours (hrs) post stimulation (10 – 100ng; Figure 1A – C). The inductions of these pro-inflammatory cytokines by CoV2-S1 were significantly greater than that induced by CoV-S1 (50ng/mL; Figure 1D – F). Higher cytokine inductions by CoV2-S1 were associated with earlier (6hr) and higher NF-κB activation (phospho (p)-p65) compared to that induced by CoV-S1 (Figure 1G).

**Figure 1.**
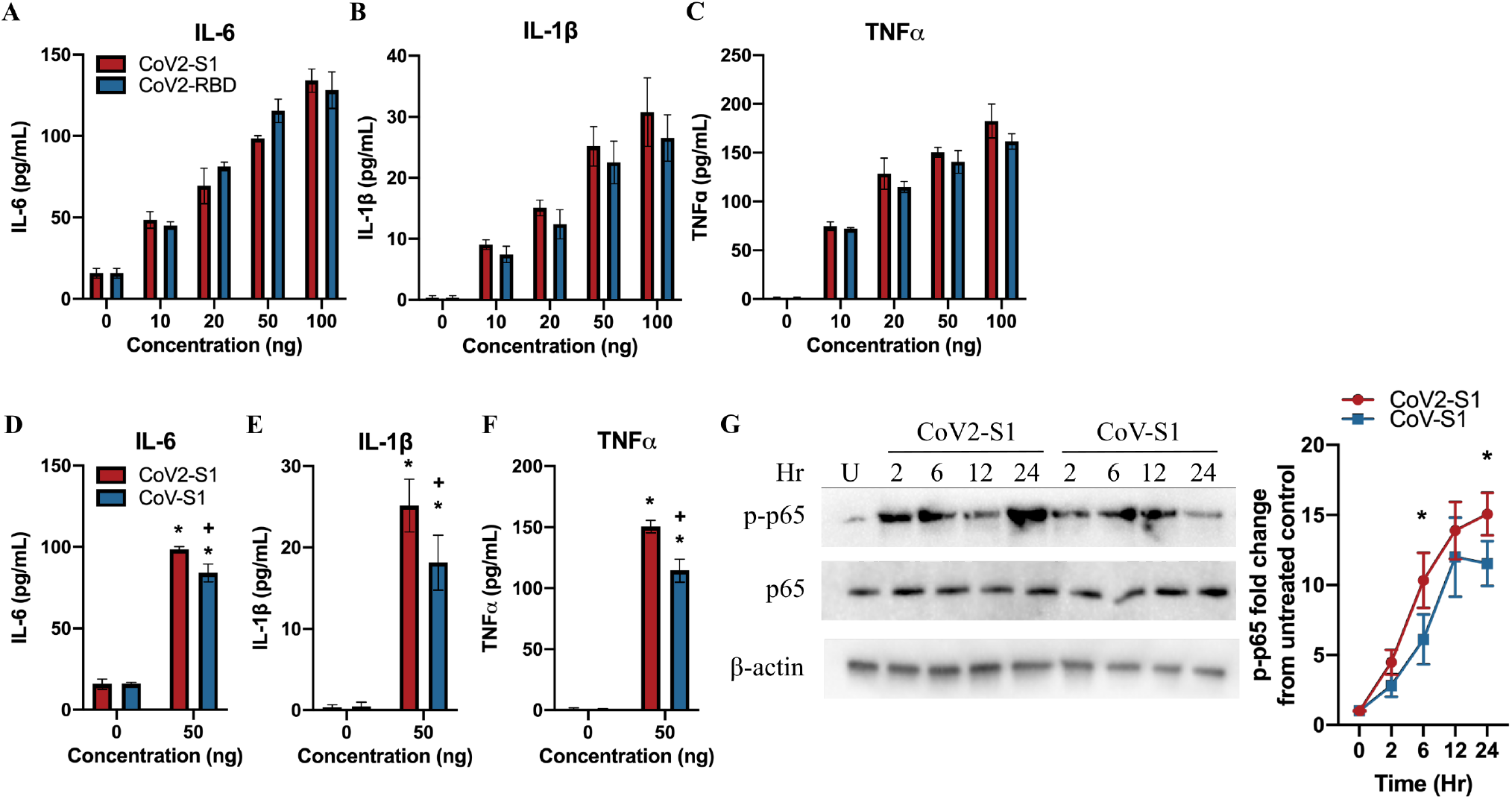
CoV2-S1 and -RBD stimulated higher productions of IL-6, IL-1β and TNF⍰ compared with CoV-S1 and -RBD. **A-C.** Protein levels of IL-6, IL-1β and TNF⍰ stimulated by SARS-CoV-2 spike subunit 1 (CoV2-S1) and receptor binding domain (RBD) 24hrs post stimulation. **D-F.** Stimulation and comparison of IL-6, IL-1β and TNF⍰ by CoV2-S1 and CoV-S1. **G.** Immunoblot (left) and densitometry (right) of induction kinetics of phospho(p)-p65 at 2, 6, 12, and 24hr post stimulation by CoV2-S1 and CoV-S1 compared with CoV2-S1 stimulated but untreated control (U). N = 6 showing mean ± SEM. Immunoblots shown are representative of three independent experiments and β-actin was detected to show equal protein input (lower panel). * *P* ≤0.05 versus untreated control.

### CoV2-S1-mediated NF-κB activation is dependent on S1-ACE2 interaction

Immunoprecipitation using anti-His antibody demonstrates that CoV2-S1 / -RBD binding to ACE2 30 minutes post stimulation (Figure 2A). To further assess if CoV2-S1 and ACE2 interaction is required for NF-κB activation, we have designed six antagonistic peptides of varying lengths (AP-1 – 6; 8 – 15 amino acid residues) that were N-terminally biotinylated (Figure 2B). These peptides were designed based on the contact residues on ACE2 in an attempt to inhibit CoV2-S1-ACE2 interaction, including Y449, Y453, L455, F486, N487, Y489, Q493, Q498, T500, N501, G502, and Y505 on CoV2-S1 (Figure 2B; Table 1) (Li et al., 2005; Wang et al., 2020a).

**Figure 2.**
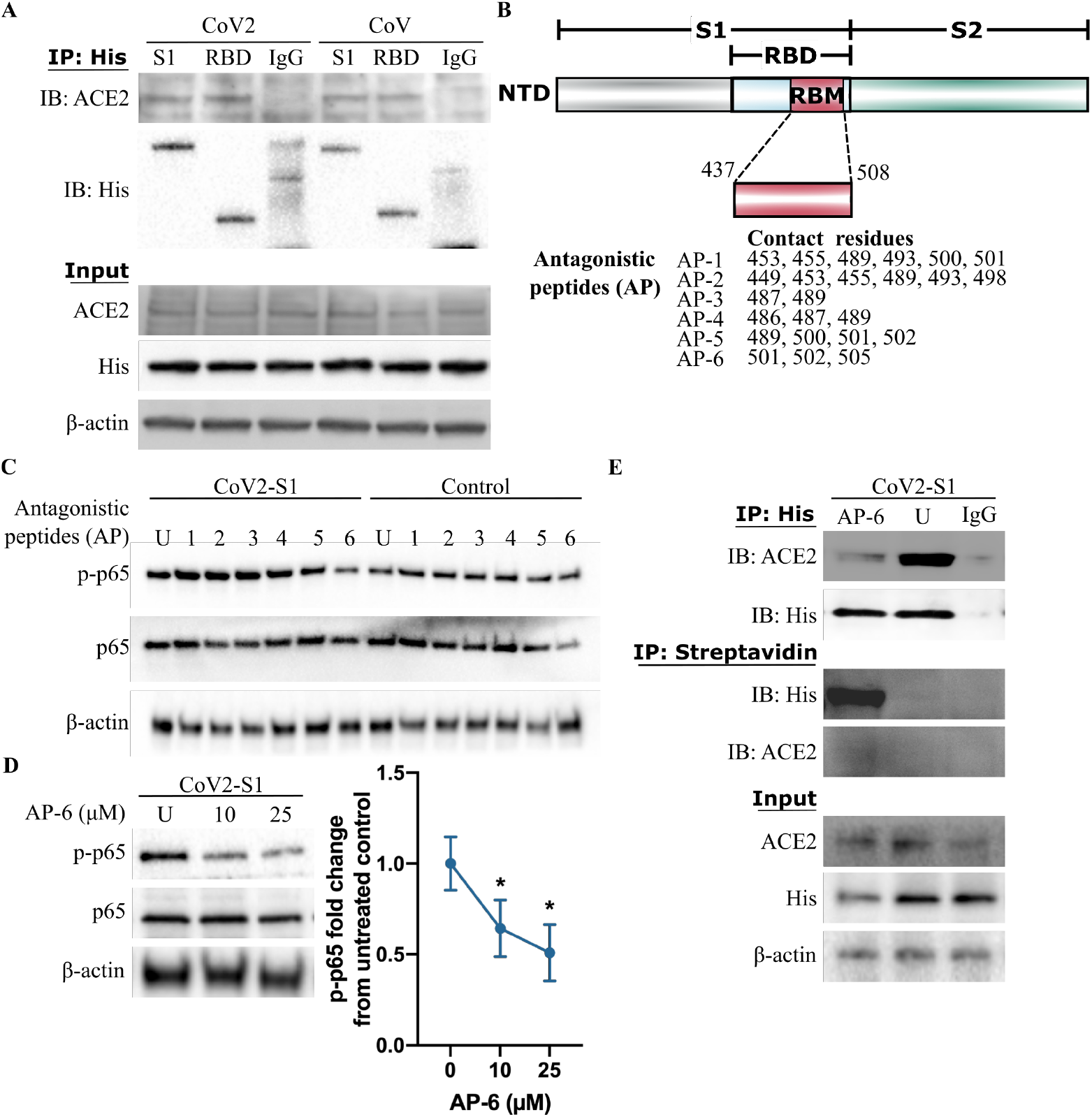
CoV2-S1-mediated NF-κB activation requires ACE2 interaction and can be inhibited by antagonistic peptides. **A.** Immunoblot of ACE2 after immunoprecipitation using anti-His antibody 30 minutes post CoV2-S1/-RBD or CoV-2-S1/-RBD stimulation. **B.** Schematic representation of CoV2-S1 protein and peptide design coverage. **C.** Immunoblot of p-p65 and p65 24hrs post CoV2-S1 stimulation and CoV2-S1 antagonistic peptides (AP-1 – 6) compared with untreated control (U). **D.** Immunoblot (left) and densitometry (right) of p-p65 and p65 at 24hrs post CoV2-S1 and AP-6 (10, 25μM) treatment compared with CoV2-S1 stimulated but untreated control (U). **E.** Immunoblot of ACE2 and His following immunoprecipitation using anti-His antibody 30 minutes post CoV2-S1/AP-6 treatment. Immunoblot of His and ACE2 following immunoprecipitation using streptavidin-coupled Dynabeads 30 mintures post CoV2-S1/AP-6 treatment. N = 3 showing mean ± SEM. Immunoblots shown are representative of three independent experiments and β-actin was detected to show equal protein input (lower panel). * *P* ≤0.05 versus untreated control.

We first screened for abilities of these peptides in reducing p65 phosphorylation by CoV2-S1.

Treatment of BCi-NS1.1 with AP-6, but not AP-1 – 5, reduced p65 phosphorylation levels induced by CoV2-S1 (Figure 2c; 10μM). AP-6 mediated reduction in p65 phosphorylation occurred in a dose-dependent manner (10 – 25μM) with minimal cell death (Figure S1A). Higher dose (50μM) resulted in increased cell death (Figure S1A). We then assessed if AP-6 inhibits S1-ACE2 interaction. In the AP-6 treated group, immunoprecipitation using anti-His antibody also shows reduced CoV2-S1 interaction with ACE2 compared with CoV2-S1-stimulated and non-AP-6-treated group (U; Figure 2E). Conversely, immunoprecipitation using streptavidin diminished interactions between CoV2-S1 and ACE2 in AP-6 group and not in the non-AP-6-treated group. AP-6 also reduced CoV-S1 and ACE2 interaction, indicating the inhibition potentials of AP-6 across small differences in amino acid residue sequences between CoV2-S1 and CoV-S1 RBM (Figure S1B – C).

This indicates that CoV2-S1 elicits higher inflammatory responses than CoV-S1, and that the CoV2-RBD alone is sufficient for inducing this response. CoV2-S1-ACE2 interaction is required for this heightened inflammatory response.

### CoV2-S1-ACE2 facilitates NF-κB activation *via* MAP kinase signalling

We investigated if S1-mediated production of pro-inflammatory cytokines requires common adaptor proteins MyD88 or TRIF. Knockdown of MyD88 or TRIF inhibited S1-mediated p65 phosphorylation (Figure 3A), indicating MyD88 and TRIF as important signalling adaptors in S1-driven p65 activation. However, immunoprecipitation using anti-His antibody did not pull down MyD88 or TRIF (Figure 3B) 2hrs post stimulation. This indicates an intermediate signalling factor is involved between ACE2 and MyD88/TRIF.

**Figure 3.**
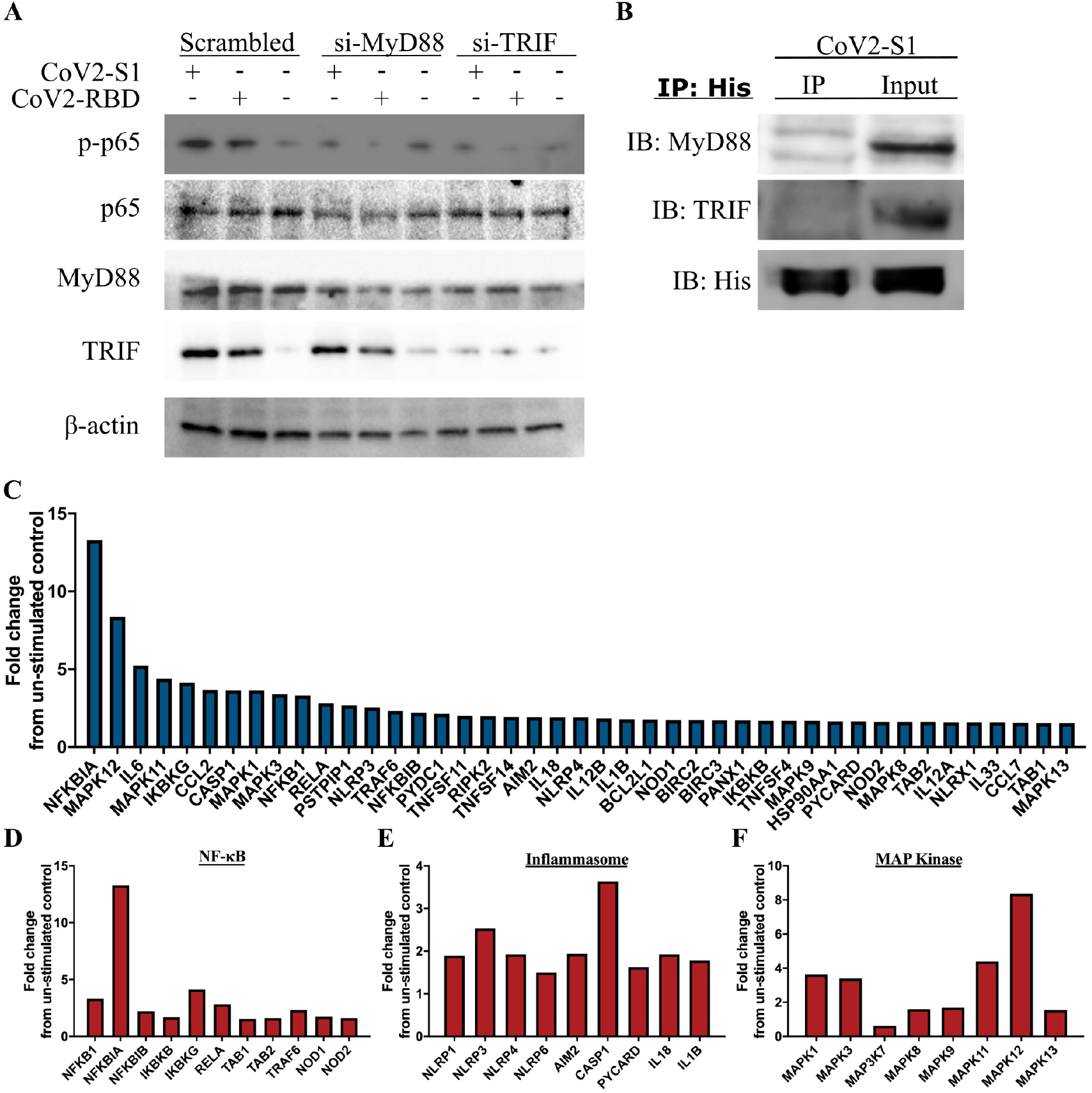
CoV2-S1 stimulates NF-κB activation via MyD88/TRIF adaptors and MAP kinase pathway. **A.** Immunoblots of p-p65 phosphorylation 24hrs post CoV2-S1 stimulation in MyD88 and TRIF silenced BCi-NS1.1. **B.** Immunoblots of MyD88 and TRIF after immunoprecipitation using anti-His antibody 2 hours post stimulation. **C-F.** CoV2-S1-stimulated BCi-NS1.1 cells (24hrs) were subjected to RT^2^ Profiler™ Human Inflammasone PCR array with highly up-regulated genes involved in NF-κB, Inflammasome, and MAP kinase pathways. N = 3 for Figure 3A-B. Immunoblots shown are representative of three independent experiments and β-actin was detected to show equal protein input (lower panel).

MyD88 and TRIF activates multiple pathways that converge at NF-κB. To determine how CoV2-S1 up-regulates NF-κB activity, we used RT^2^ Profiler™ Human Inflammasone PCR array and investigated intracellular signalling pathways involved in NF-κB activation (Figure 3C). CoV2-S1 stimulation in BCi-NS1.1 increased expression of genes involved in NF-κB (*NFKB1, NFKBIA, NFKBIB, RELA, TAB1*) and inflammasome signalling (*NLRP1/3/4/6, AIM2, CASP1, PYCARD, IL1B, and IL18*), but also several factors involved in MAP kinase pathway (*MAPK1/3/3K7/8/9/11/12/13*) (Figure 3D – F; Figure S2).

MAP kinase pathways have been shown to be involved in NF-κB activations (Bergmann et al., 1998; Madrid et al., 2001; Wang et al., 2019; Wesselborg et al., 1997), and consistent with the PCR array. Here CoV2-S1 stimulation in BCi-NS1.1 cells also induced heightened activation of MAP kinase p38 (12 – 24hrs), Erk (2 and 24hrs), and Jnk (24hrs), compared to inductions by CoV-S1 (Figure 4A). Knockdown of p38, Erk or Jnk expressions by siRNAs all resulted in a significant reduction in phosphorylation of p65 following CoV2-S1 treatment (Figure 4B), indicating that the MAP kinase is an important contributor to CoV2-S1 - mediated NF-κB activation.

**Figure 4.**
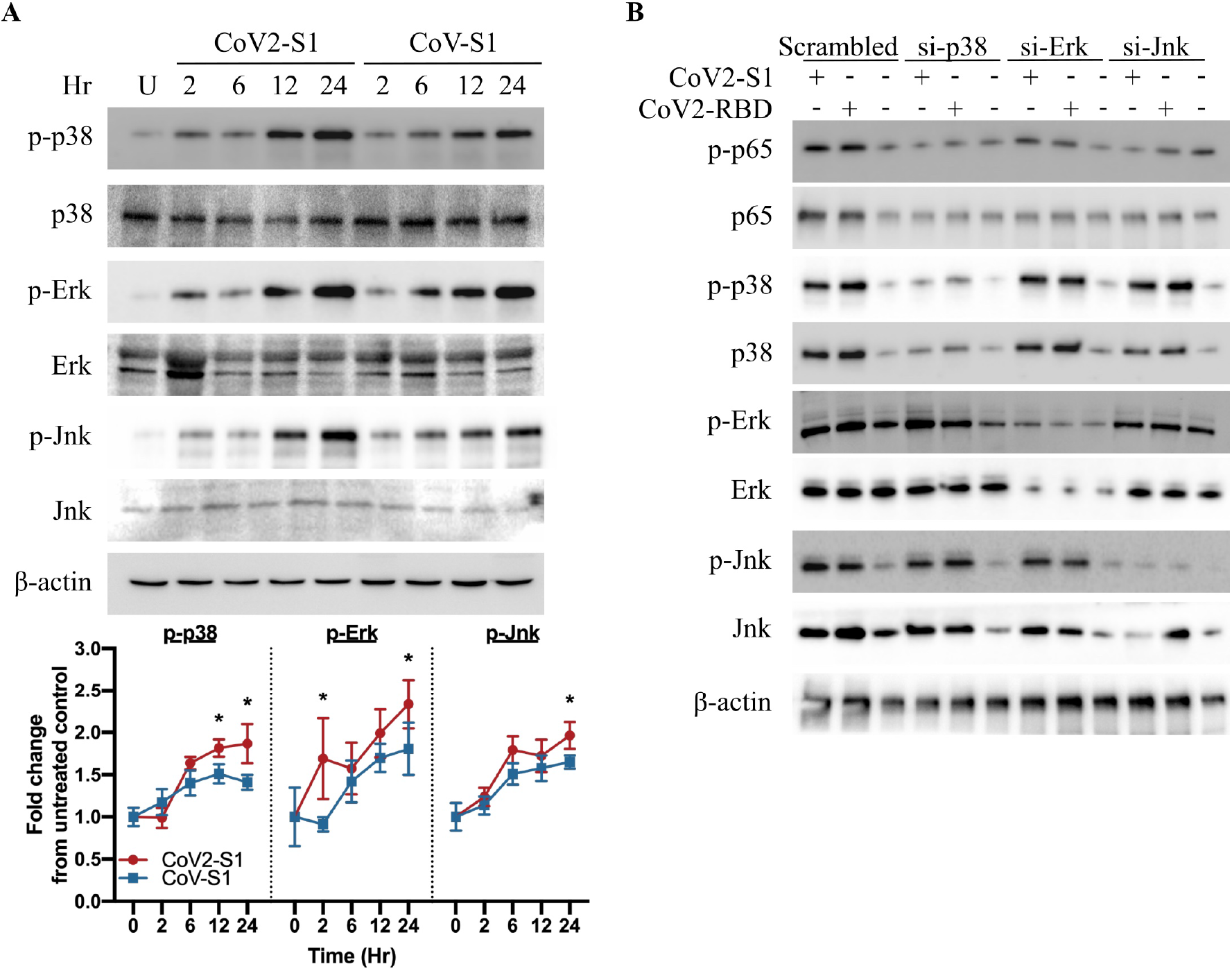
CoV2-S1 mediates NF-κB activation *via* MAP kinase pathways. **A.** Immunoblot (top) and densitometry (bottom) of induction kinetics of p-p38, p38, p-Erk, Erk, p-Jnk and Jnk at 2, 6, 12, and 24hr post stimulation by CoV2-S1 and CoV-S1 compared with untreated control (U). **B.** Immunoblots of p-p65, p65, p-p38, p38, p-Erk, Erk, p-Jnk and Jnk 24hrs post CoV2-S1 or -RBD stimulation in p38, Erk or Jnk silenced BCi-NS1.1. N = 3. Immunoblots shown are representative of three independent experiments and β-actin was detected to show equal protein input (lower panel). * *P* ≤0.05 versus untreated control.

### CoV2-S1 stimulates rapid ER stress and UPR that activate NF-κB *via* UPR-MAP kinase crosstalk

CoV2-S1 induced early activation of Erk, which is a kinase utilised by not only MAP kinase but also ER stress and UPR. ER stress has been shown to be induced by the CoV spike protein (Versteeg et al., 2007). ER-UPR features three main pathways, PERK-Erk-CHOP pathway that modulates apoptosis, ATF6 that regulates protein folding, and the IRE1⍰ – TRAF2 pathway that promotes NF-κB and p38 activation (Pathinayake et al., 2018).

CoV2-S1 and -RBD resulted in an early increase of IRE1⍰ and PERK activations at 2 and 6hrs, respectively, and this was sustained to 24hr post stimulation. Furthermore, IRE1⍰ and PERK activations were higher compared with CoV-S1 and -RBD (Figure 5A). ATF6 activation was equivalent for CoV2-S1 and CoV-S1. As IRE1⍰ activation was induced earlier (2hrs) by CoV2-S1 than MAP kinases, we then investigated if increased ER-UPR by CoV2-S1 leads to heightened MAP kinase activities by siRNAs.

**Figure 5.**
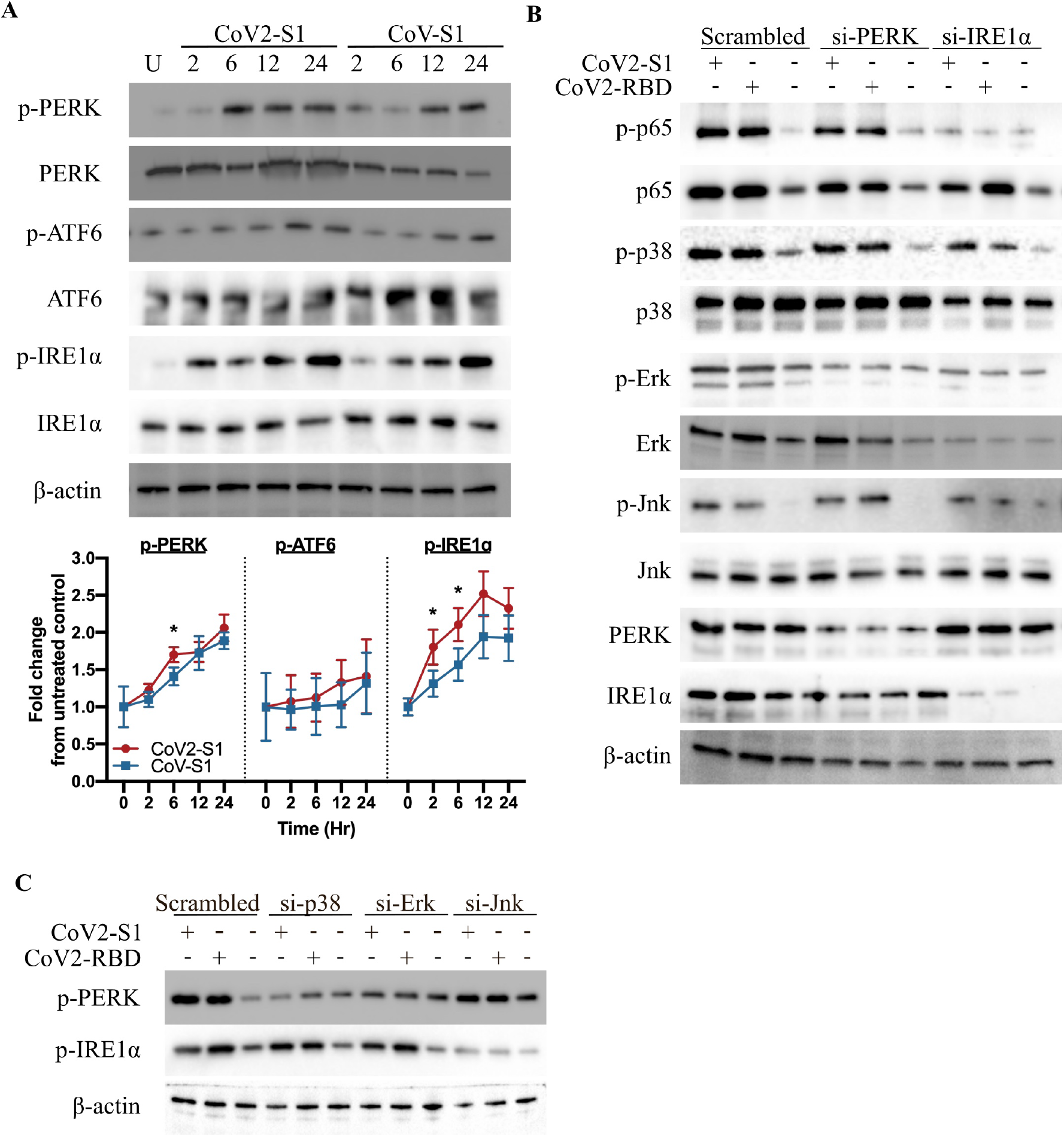
CoV2-S1 induces rapid ER-UPR that activate NF-κB *via* ER-MAP kinase crosstalk. **A.** Immunoblot (top) and densitometry (bottom) of induction kinetics of p-IRE1⍰, IRE1⍰, p-ATF6, ATF6, p-PERK and PERK at 2, 6, 12, and 24hr post stimulation by CoV2-S1 and CoV-S1 compared with untreated control (U). **B.** Immunoblots of p-p65, p65, p-p38, p38, pErk, Erk, p-Jnk, Jnk, IRE1⍰ and PERK 24hrs post CoV2-S1 or -RBD stimulation in PERK and IRE1⍰ silenced BCi-NS1.1. **C.** Immunoblots of p-PERK and p-IRE1⍰ post CoV2-S1 or -RBD stimulation in p38, Erk or Jnk silenced BCi-NS1.1. N = 3. Immunoblots shown are representative of three independent experiments and β-actin was detected to show equal protein input (lower panel). * *P* ≤0.05 versus untreated control.

Knockdown of PERK resulted in reduced phosphorylation levels of Erk, but had minimal effect on p65, p38, and Jnk activation (Figure 5B). In contrast, reduction of IRE1□ expression decreased p65, p38, and Erk phosphorylation/expression. This indicates that IRE1⍰, and not PERK and ATF6 pathway, is involved in p65 activation.

We also assessed if MAP kinase modulates ER-UPR. While knockdown of either p38, Erk, or Jnk decreased p65 phosphorylation (Figure 4B), reduction of p38 or Erk gene expression led to decreased PERK phosphorylation but not IRE1⍰ activation (Figure 5C). Jnk knockdown led to reduced IRE1⍰ phosphorylation and had no effect on PERK activation. This demonstrates CoV2-S1 promotes NF-κB activation *via* the ER-UPR (IRE1⍰/PERK) and MAP kinase pathway.

### CoV2-S1 induces markers of acute bronchial epithelial injury

Acute lung injury is a feature observed with severe COVID-19 (Huang et al., 2020; Lai et al., 2017), and markers associated with epithelial damage IL-1β, TNF⍰, and CCL2 were all markedly increased by CoV2-S1 in our targeted PCR array (Figure 3C), and at protein levels (Figure 1A). CoV2-S1 also significantly up-regulated protein productions of CCL2 in BCi-NS 1.1 cells (Figure 6A). To further investigate if CoV2-S1 induces epithelial damage, we stimulated differentiated primary bronchial epithelial cells (pBECs) cultured at air-liquid interface (ALI) with CoV2-S1 (50 and 100ng). Stimulation resulted in a significant reduction in transepithelial electrical resistance (TEER; Figure 6B), demonstrating an increased epithelial permeability and tight junction disruption caused by CoV2-S1. This is accompanied with increased protein production of epithelial damage factors IL-1β, TNF⍰, and CCL2, in a CoV2-S1 dose-dependent manner (Figure 6C – E).

**Figure 6.**
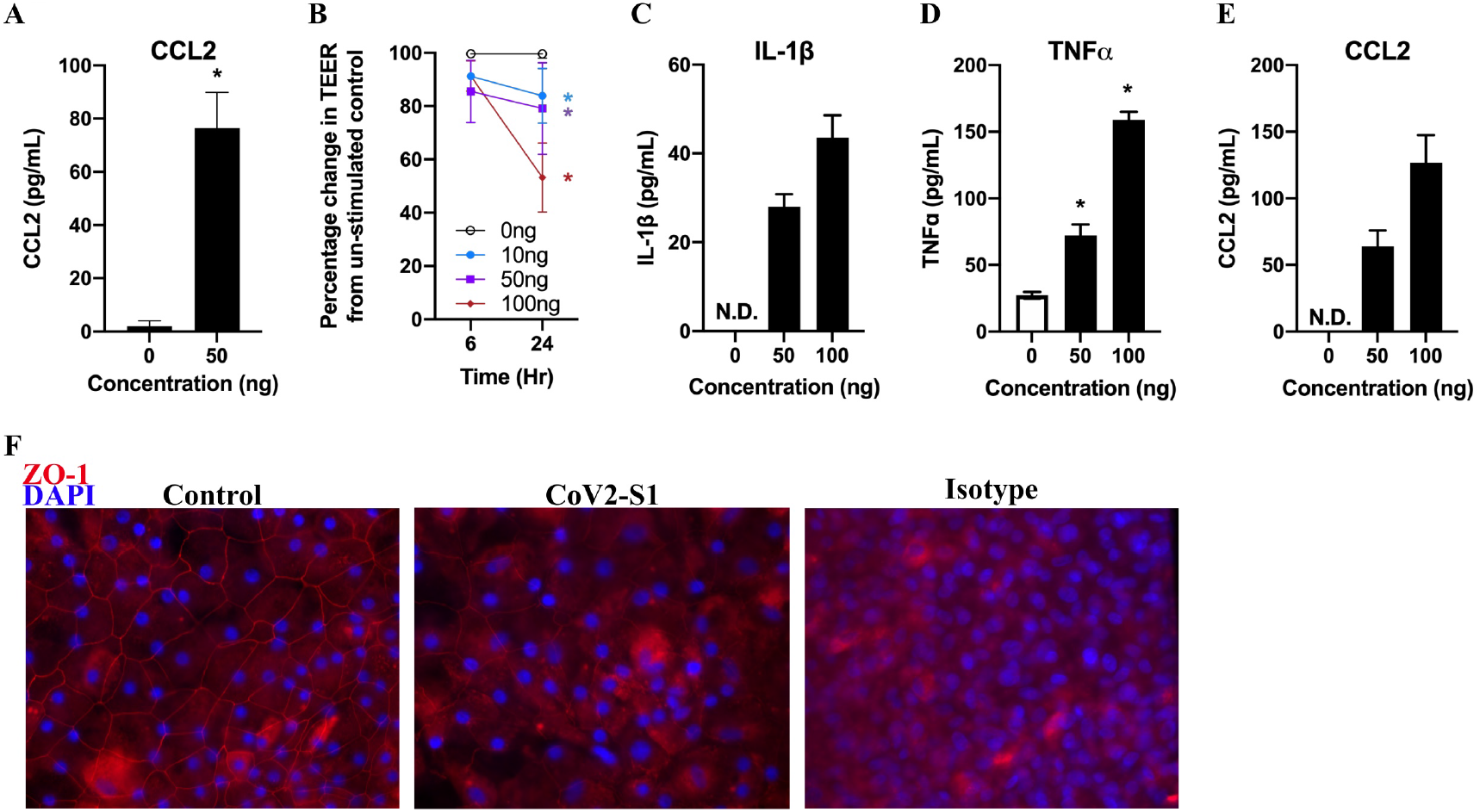
CoV2-S1 promotes epithelial damage. **A.** CCL2 protein production at 24hrs post CoV2-S1 stimulation in BCi-NS1.1. **B.** Percentage changes in transepithelial electrical resistance (TEER) in differentiated primary bronchial epithelial cells (pBECs) cultured at air-liquid interface (ALI) 24hrs post CoV2-S1 stimulation. **C-E.** Protein levels of IL-1β, TNF⍰, and CCL2 at 24hrs post CoV2-S1 stimulation in pBECs-ALI. N = 3. * *P* ≤0.05 versus untreated control. **F.** Immunofluorescent images of ZO-1 labelling. N = 2.

Immunofluorescent staining of a tight junction protein zonula occludens-1 (ZO-1) showed strong localisation at the cell borders in the non-stimulated controls, whereas CoV2-S1 led to partial disappearance of ZO-1 at the cell borders (Figure 6F). This indicates that CoV2-S1 may also disrupt epithelial barrier function.

### CoV2-S1-antagonistic AP-6 and FDA-approved ER stress and MAP kinase inhibitors ameliorate CoV2-mediated inflammatory response

CoV2-S1 induced NF-κB activation and subsequent production of pro-inflammatory cytokines was dependent on ACE2 interaction and early ER-UPR and MAP kinase activities. We therefore assessed whether CoV2-S1-mediated inflammation could be reduced by our CoV2-S1 inhibitory peptide AP-6, or with existing FDA-approved pharmacological inhibitors, that target ER stress (4-PBA) or MAP kinase (trametinib and ulixertinib).

Treatment with AP-6 led to a significant decrease in CoV2-S1-mediated phosphorylation of p65 (Figure 2C) and production of IL-6, IL-1β, TNF⍰, but not CCL2 (Figure 7A – D) in a dose-dependent manner. Similarly, the ER stress inhibitor 4-PBA and both MAP kinase inhibitors led to decreased activation of p65 (Figure S3) and expression of these pro-inflammatory and epithelial injury cytokines (Figure 7E – L). These drugs had no effect on non-CoV2-S1-stimulated controls (Figure S4). CoV2-S1-induced reduction in barrier integrity and ZO-1 was prevented *via* treatment with AP-6 and FDA-approved inhibitors, suggesting maintained barrier function (Figure S5A – B). This strongly indicates that AP-6 could serve as a proof-of-concept therapeutic peptides against SARS-CoV-2 and also that ER stress / MAP kinase inhibitors could be used to reduce inflammation and lung injury caused by CoV2-S1.

**Figure 7.**
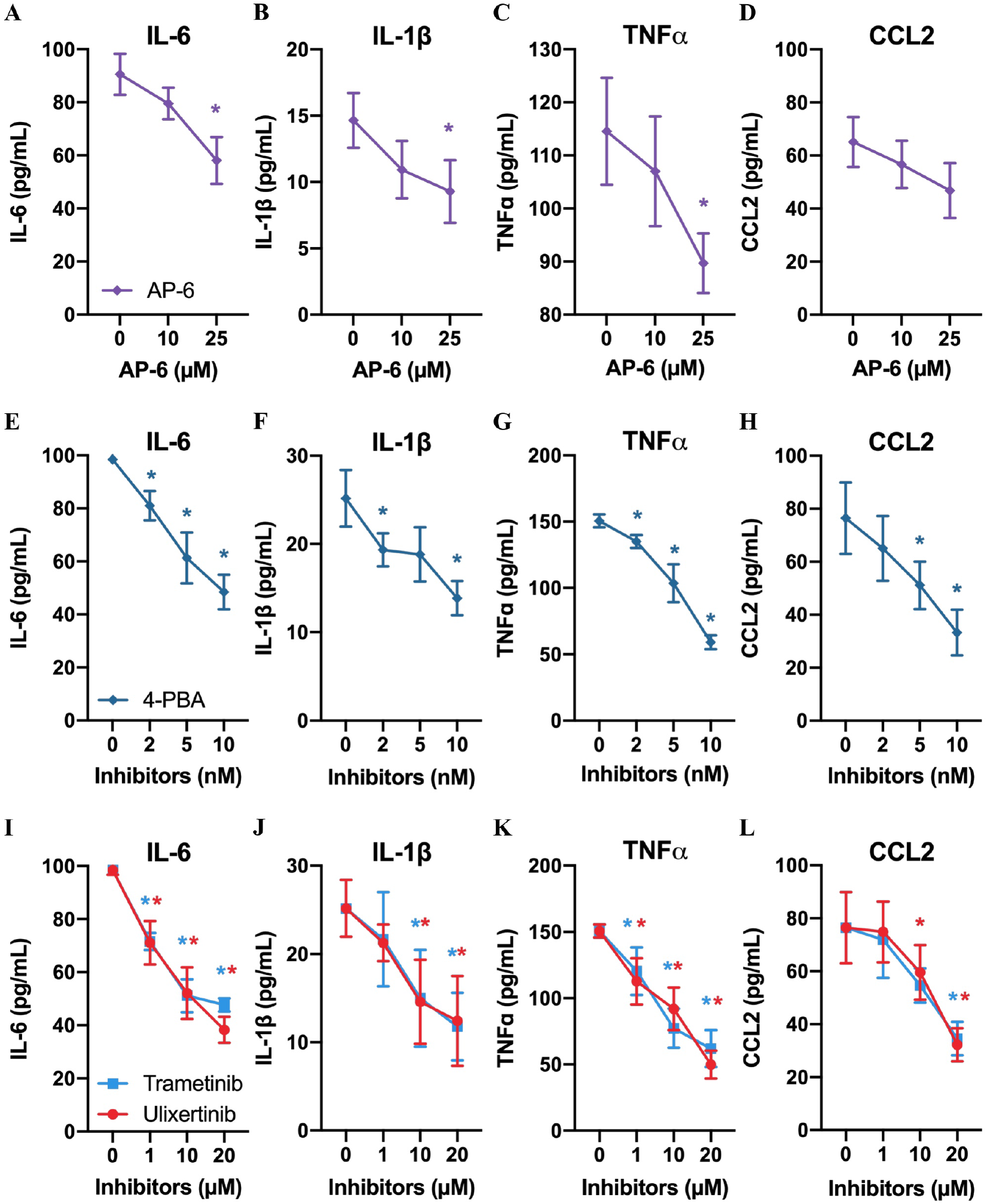
CoV2-S1 antagonistic peptide AP-6 and FDA-approved ER stress and MAP kinase inhibitors suppress CoV2-S1-mediated production of pro-inflammatory cytokines. **A-D.** Protein levels of IL-6, IL-1β and TNF⍰ at 24hrs post stimulation by CoV2-S1 and treatment with antagonistic peptide AP-6, **E-H.** 4-PBA, and **I-L.** Trametinib, and Ulixertinib. N = 3. * *P* ≤0.05 versus untreated control.

## Discussion

CoV Spike protein and host ACE2 interaction is a critical first step to viral replication and diseases. Here we demonstrate that CoV2 S1 subunit and RBD induces early ER-UPR and MAP kinase activations, leading to hyper-inflammatory responses. Our results indicate that this inflammatory storm, and downstream consequences are inhibited by S1-inhibitory peptides and existing FDA-approved ER stress and MAP kinase inhibitors (Figure 8).

**Figure 8.**
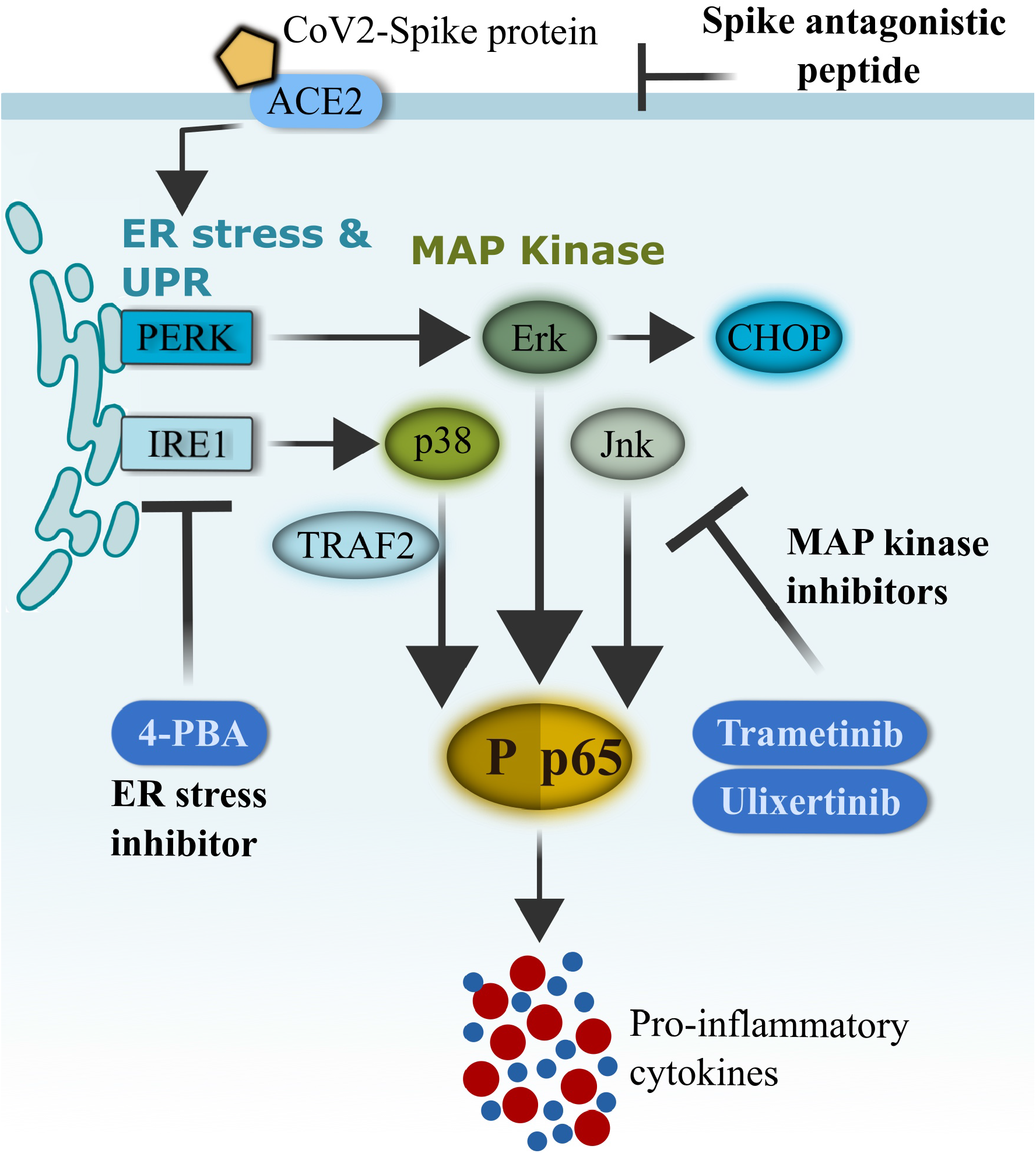
Schematics of CoV2-S1-mediated inflammation via ER-UPR and MAP kinase pathways. SARS-CoV-2 (CoV2) spike protein binds with ACE2 on the surface of human bronchial epithelial cells and rapidly facilitates the induction of ER stress and unfolded protein response (UPR). Activation of UPR (PERK and IRE1□) promotes the activation of MAP kinases, and the two pathways synergistically drive the activation of NF-κB and production of pro-inflammatory cytokines. FDA-approved ER-UPR inhibitor 4-phenylburic acid (4-PBA) and MAP kinase inhibitors (trametinib and ulixertinib) suppressed CoV2-S1-induced ER stress and MAP kinase activities, resulting in reduced NF-κB-mediated expression of pro-inflammatory cytokines.

COVID-19 has been shown to be associated with increased plasma levels of pro-inflammatory cytokines including IL-6 and TNF□ (Huang et al., 2020; McGonagle et al., 2020; Mehta et al., 2020). SARS-CoV-2 S1 and RBD alone induced heightened levels of IL-6, TNF□ and IL-1β, and the productions of which were higher compared with CoV-S1. This induction occurs in an ACE2-dependent manner, and the higher affinity of CoV2-S1 towards ACE2 may have contributed to this elevated inflammatory response (Wang et al., 2020a). CoV2-S1-mediated NF-κB activation is also dependent on common TLR adaptor proteins MyD88 and TRIF, although we could not detect interactions between ACE2 and these adaptor proteins. It is possible that CoV2-S1-ACE2 interaction triggers MyD88 and TRIF activations *via* other signalling factors that then result in NF-κB activation.

Our data strong indicates that inflammation could be triggered by CoV2-S1 even before viral replication occurs or in the absence of viral replication. As CoV2-S1 is present throughout viral replication cycles and infection, our data demonstrate that spike proteins are likely to be a major contributor to inflammation. The constant presence of S1 is consistent with the high viral load and a long virus-shedding period observed in patients with severe COVID-19 (Liu et al., 2020). Furthermore, CoV2 spike protein has an estimated half-life of 30 hours in mammalian systems (*in silico* analysis by ExPASy ProtParam) and consists of amino acid residues with long half-lives (leucine (8%), threonine (7.6%), valine (7.6%), alanine (6.2%), glycine (6.4%), and proline (4.6%)). This may suggest persistent presence of CoV2 spike protein from live and dead viruses that plays major role in triggering the inflammatory storm in COVID-19. Importantly, our antagonistic peptide inhibited this binding event and reduced the production of pro-inflammatory cytokines. While peptide sequence optimisations are required to further increase effectiveness and stability, this highlights the potentials of using S1 antagonistic peptides as neutralising molecules against SARS-CoV-2.

The ER is mainly involved in protein folding, trafficking and post-translational modification of secreted and transmembrane proteins (Pathinayake et al., 2018). Viral infections and inflammatory cytokines result in high ratio of misfolded or unfolded proteins in the ER, leading to UPR (Oslowski and Urano, 2011). This adaptive mechanism triggers a range of different innate immune responses *via* different UPR pathways; PERK-Erk-CHOP pathway that halts protein translation (Harding et al., 2000); ATF6 that facilitates protein folding as well as NF-κB activation (Shen et al., 2005); and IRE1□ promotes NF-κB phosphorylation and inflammation (Urano et al., 2000). Here we showed that both ER-UPR and MAP kinases modulate CoV2-S1-induced NF-κB activations.

CoV2-S1 caused early activation of ER-UPR IRE1⍰ and PERK-Erk, leading to MAP kinases phosphorylations, which then facilitated NF-κB-mediated production of pro-inflammatory cytokines. Although IRE1⍰ activation occurred earlier than other ER-UPR and MAP kinase factors, it is likely that CoV2-S1 can promote NF-κB activation *via* ER-UPR and MAP kinase both independently and cooperatively (Bergmann et al., 1998; Madrid et al., 2001; Wesselborg et al., 1997) (Harding et al., 2000; Urano et al., 2000). Our result is consistent with previous reports that showed ER stress and MAP kinase activation by SARS-CoV infection (Lee et al., 2004; Mizutani et al., 2004; Versteeg et al., 2007), we cannot rule out the possibilities of other mechanisms of ER-UPR and MAP kinase activations by CoV2-S1. Furthermore, during SARS-CoV-2 infection, viral RNAs will also trigger inflammatory responses *via* multiple pattern recognition receptors including TLRs and RLRs. This together with spike proteins may instigate more exaggerated inflammatory responses during SARS-CoV-2 infection.

Acute lung injury is a feature of severe COVID-19 (Huang et al., 2020; McGonagle et al., 2020; Mehta et al., 2020). Surprisingly we found that CoV2-S1 and -RBD alone was also sufficient to reduce epithelial barrier function through the release of IL-1β, TNF⍰, and CCL2. Pro-inflammatory cytokines IL-1β and TNF⍰ have been shown to induce epithelial damage by further promoting p38 and NF-κB activation (Al-Sadi et al., 2013; Kimura et al., 2013; Kimura et al., 2009; Kolb et al., 2001; Sheridan et al., 1997). CCL2 is a chemokine that has been shown to be transcriptionally driven by NF-κB and is typically released by injured tissues and attracts macrophages to the site of infection and inflammation (Kavandi et al., 2012; Lai et al., 2017).

While CoV2-S1 increased membrane permeability that was consistent with small loss of ZO-1 localisation at cell-cell junctions, it is likely that increased production of these inflammatory and injury-related factors from epithelial cells recruit macrophages to the site of infection, which then promote excessive tissue damage. These inflammatory and injury-stimulated factors as well as macrophages have been reported to be highly elevated in those with severe COVID-19 (Huang et al., 2020; McGonagle et al., 2020; Mehta et al., 2020), further indicating that this “inflammatory injury” can be driven by CoV2-S1 in severe COVID-19, and may be the first step to ARDS.

Increased ER-UPR and MAP kinase activities as well as pro-inflammatory responses induced by CoV2-S1 could be substantially reduced by FDA-approved ER stress inhibitor 4-PBA and MAP kinase inhibitors trametinib and ulixertinib. 4-PBA is a chemical chaperone currently used for treatment of urea cycle disorder (Lichter-Konecki et al., 2011), and has been used in clinical trials for diabetes (Ozcan et al., 2006), cystic fibrosis (Ozcan et al., 2006), sickle cell disease (Collins et al., 1995), and neurodegenerative diseases (Mimori et al., 2012). Trametinib is a MAP kinase inhibitor used for melanoma (Hoffner and Benchich, 2018), and ulixertinib is a highly potent, selective, reversible, ERK1/2 inhibitor used as cancer treatment (Sullivan et al., 2018). Re-purposing these drugs that has a well-documented safety profile in humans could expedite rapid deployment of these drugs for severe COVID-19.

Collectively, CoV2-S1 induced heightened production of inflammatory cytokines that are primarily driven by MAP kinase and ER stress cross-talks. CoV2-S1 AP-6 demonstrates the feasibility of this proof-of-concept CoV-specific antiviral strategy. While antiviral drugs and vaccines are being developed and assessed, existing FDA-approved ER stress and MAP kinase inhibitors could be immediately deployed in clinical trials as a potential treatment options for those with severe COVID-19.

## Acknowledgments

This study is funded by University of Newcastle Faculty Pilot Grant (1032380) and John Hunter Hospital Charitable Trust (G1700465).

The authors have declared that no conflict of interest exists.

## Author Contributions

Conceptualization, A. C-Y. H.; Methodology, A. C-Y. H., G. W., A. T. R., P. C. V., P. S.P.; Investigation, A. C-Y. H., G. W., K. D., A. T. R., P. C. V.; Validation, G. W., A. T. R., P. C. V., P. S.P., K. D.; Writing – Original and finalised version, A. C-Y. H.; Writing – Review & Editing, A. C-Y. H., G. W., A. T. R., P. C. V., P. S.P., K. D., J. C. H., F. W., P. A. W.; Funding Acquisition, A. C-Y. H.; F. W.

## STAR Methods

### Experimental model and subject details

#### Cell line

BCi-NS1.1 cells were obtained from Prof. Ronald Crystal Laboratory at Weill Cornell Medical College, and Memorial Sloan-Kettering Cancer Center, New York, NY, USA) (Walters et al., 2013). The cells were cultured in hormone supplemented Bronchial Epithelial Cell Growth Media (BEGM; Lonza, Switzerland) supplemented with 50U/mL penicillin and streptomycin (Hayman et al., 2019; Hu et al., 2019; Kedzierski et al., 2017).

#### Human subject recruitment for pBECs

Five healthy control subjects were recruited for bronchoscopy. Healthy non-smoking controls with no evidence of airflow obstruction, bronchial hyper-responsiveness to hypertonic saline challenge, or chronic respiratory symptoms were also recruited. Clinical history, examination and spirometry were performed on all individuals, whom were also questioned about the previous severity of cold symptoms. At the time of recruitment none of the subjects had symptoms of acute respiratory tract infections for the preceding four weeks and did not have a diagnosis of lung cancer.

All subjects gave written informed consent. All procedures were performed according to approval from The University of Newcastle Human Ethics Committees

#### Differentiation of primary bronchial epithelial cells (pBECs) at air-liquid-interface (ALI)

Human pBECs were obtained by endobronchial brushing and research bronchoscopy in accordance with standard guidelines. pBECs were cultured in BEGM in polycarbonate tissue culture flasks as previously described (Hsu et al., 2011; Hsu et al., 2017; Hsu et al., 2012; Hsu et al., 2016; Hsu et al., 2015; Kedzierski et al., 2017; Parsons et al., 2014; Vanders et al., 2019), and were then cultured on polyester membrane transwells (12mm diameter, 0.4μM pore size, Corning, USA) under submerged condition in ALI initial media (31.25mL low glucose DMEM and BEGM, 1μL of 1mM All-trans retinoic acid, 4μL of 25μg/mL recombinant human epidermal growth factor (rhEGF), 62.5μL hydrocortisone, bovine insulin, epinephrine, transferrin, 80μM ethanolamine, 0.3mM MgCl2, 0.4mM MgSO4, 0.5mg/mL BSA, and 250μL bovine pituitary extract, supplemented with penicillin/streptomycin and amphotericin B). When cells become fully confluent, pBECs were air-lifted by removing apical media and changing basal media into ALI final media (as described above for ALI initial media but with 0.5ng/mL of rhEGF). Transepithelial resistance (TEER) was measured every seven days using a EVOM2 Epithelial Voltohmmeter (World Precision Instruments, USA). The basal media was replaced with fresh ALI final media every second day. pBECs were cultured at ALI for 25 days and then used for experiments. All cells were cultured and maintained at 37°C / 5% CO_2_.

### Method details

#### Spike protein subunit 1 (S1) and receptor binding domain (RBD) stimulation

For cells cultured in submerged and at ALI conditions, his-tagged S1 and RBD (Sino Biological Inc.) was diluted in BEBM minimal media (Lonza, Switzerland) and added to the cells. For pBECs at ALI, S1 and RBD was added to the apical side.

#### CoV2-S1 antagonistic peptides

Six peptides of varying lengths (8 – 15 amino acid residues) were designed based on the amino acid residues critical in ACE2 binding within the receptor binding motif (RBM) of the receptor binding domain (RBD). The peptides were biotinylated at the amino-terminus with carboxy-terminal amidation. The peptides were synthesised by GenScript Biotech Corp. The peptides were added to the BCi-NS1.1 at 10 or 25μM, or added to pBECs-ALI (25μM) at the apical side.

#### Drugs

ER stress inhibitor 4-phenylburic acid (4-PBA) was purchased from Sigma-Aldrich, and resuspended in H_2_O and diluted in culture media. MAP kinase inhibitors trametinib (GSK1120212) and ulixertinib (BVD-523) were purchased from Selleck Chemicals and were resuspended and diluted in DMSO.

#### Cell viability

Cell viability was measured using PE Annexin V Apoptosis Detection kit I (Becton Dickinson) according to manufacturer’s instruction. Cells were stained with annexin V-PE and vital dye 7-amino-actinomycin (7-AAD), and then analyzed using a FACSCanto II (Becton Dickinson) and FACSDiva software. Viable cells were stained AxV negative / 7-AAD negative and expressed as percentage of total analyzed cells.

#### siRNAs

siRNAs specific to MyD88, TRIF, p38, Erk, Jnk, PERK, and IRE1 □ (Life Technologies, USA) were reverse transfected into cells using siPORT NeoFX transfection agent (Ambion, USA) according to manufacturer’s instruction. Silence Negative controls (Life Technologies, USA) were used as siRNA negative controls.

#### PCR array

RNAs from S1-stimulated cells were extracted using RNeasy Mini Kits and QIAcube (Qiagen, USA) according to manufacturer’s instruction. 200ng of RNAs were reverse transcribed to cDNA using random primers (Applied Biosystem, USA). cDNAs were then subjected to pathway-focused gene expression array using RT^2^ Profiler™ Human Inflammasome PCR array (Qiagen, USA). The raw data was analysed by the Data Analysis Center on Qiagen website (https://www.qiagen.com/mx/shop/genes-and-pathways/data-analysis-center-overview-page/).

#### Immunoblotting and immunoprecipitation

S1-stimulated cells were lysed in protease-inhibitor cocktail supplemented RIPA buffer (Roche, UK). Proteins were subjected to SDS-PAGE (Bio-Rad Laboratories, USA) and transferred onto polyvinylidene fluoride membranes (Merck-Milipore, USA). Proteins were detected using antibodies to His, MyD88, TRIF, p65, phospho-(p)-p65, p38, p-p38, Erk, p-Erk, Jnk, p-Jnk, PERK, p-PERK, IRE1□, and p-IRE1□ antibodies (All from Abcam, UK). Antibody to ACE2 was obtained from RnD Systems (USA). For immunoprecipitation, whole cell lysates (1mg) were immunoprecipitated using anti-His antibody or isotytpe antibody (Abcam, UK), streptavidin-coupled Dynabeads™ M-280 and Dynabeads™ His-tag isolation & Pulldown kit (Life Technologies, USA) according to manufacturer’s instruction. Proteins were detected using SuperSignal WestFemto Maximum Sensitivity Substrate (Thermo Fisher Scientific, USA) and visualised on a ChemiDoc MP Imaging system (Bio-Rad Laboratories, USA).

#### Immunofluorescent microscopy

Treated pBECs cultured at ALI were fixed 4% paraformaldehyde and blocked with 50mM glycine overnight, and then stained with anti-ZO-1 antibody (Thermo Fisher Scientific, USA) or rabbit isotype IgG (Abcam, UK), counter stained with DAPI (Life Technologies, USA), and viewed under a Axio Imager M2 microscope and analyzed using Zen imaging software (Zeiss) as described previously (Liu et al., 2019; Liu et al., 2016; Liu et al., 2017; Reid et al., 2020).

#### Cytometric bead array and ELISA

Human IL-6, IL-1β, and TNF□ concentrations were determined by cytometric bead array and flow cytometry (FACSCanto II flow cytometer; BD Biosciences, USA) according to the manufacturer’s instructions. Human CCL2 and TGFβ was measured by ELISA according to the manufacturer’s instructions (R&D Systems, USA).

### Quantification and statistical analysis

#### Statistical analysis

Data were analysed on GraphPad Prism 8. Statistical significance of differences was assessed using parametric Student’s two tailed t test for normally distributed data and Mann-Whitney U test for non-parametric data. Differences were considered significant when p< 0.05.

---

**Figure S1. AP-6 reduces CoV-S1 and ACE2 interaction.**

**A.** Cell viability measured by annexin-V/7-AAD staining and flow cytometry. **B.** Immunoblot of ACE2 and His following immunoprecipitation using anti-His antibody 30 minutes post CoV2-S1/AP-6 treatment. N = 3. Immunoblots shown are representative of three independent experiments. **C.** Amino acid residue sequence alignments of CoV2-S1 and CoV-S1. Blue = AP-6 contact region. Sequence alignment performed by Clustal Omega.

**Figure S2. CoV2-S1 up-regulates genes involved in inflammation, inflammasome and MAP kinase pathways.**

CoV2-S1-stimulated BCi-NS1.1 cells were subjected to RT^2^ Profiler™ Human Inflammasone PCR array with highly up-regulated genes involved in NF-κB, Inflammasome, and MAP kinase pathways. N=3.

**Figure S3. CoV2-S1 antagonistic peptides, Trametinib, and Ulixertinib had no effect on IL-6, IL-1β, TNF⍰, and CCL2 productions in non-CoV2-S1-treated cells.**

Protein levels of IL-6, IL-1β, TNF⍰, and CCL2 induced by **A-D** antagonistic peptide (AP-6), **E-H.** 4-PBA, **I-L.** trametinib, and **M-P.** ulixertinib. N = 3.

**Figure S4. FDA-approved ER stress and MAP kinase inhibitors suppress CoV2-S1-mediated NF-κB activation.**

Immunoblot (top) and densitometry (bottom) of p-p65 and p65 at 24hr post stimulation by CoV2-S1 or -RBD compared with vehicle control. Immunoblots shown are representative of three independent experiments and β-actin was detected to show equal protein input (lower panel).

**Figure S5. AP-6, 4-PBA, trametinib and ulixertinib improves epithelial permeability.**

**A.** Percentage changes in transepithelial electrical resistance (TEER) in differentiated primary bronchial epithelial cells (pBECs) cultured at air-liquid interface (ALI) at 24hrs post CoV2-S1 stimulation and treatment with either AP-6, 4-PBA, trametinib, and ulixertinib. N = 3. * *P* ≤0.05 versus non-CoV2-S1-stimulated control, + versus CoV2-S1-stimulated group. **B.** Immunofluorescent images of ZO-1 staining. N = 2.

