## Supplemental figures for "SARS-CoV-2 Spike protein promotes hyper-inflammatory response that can be ameliorated by Spike-antagonistic peptide and FDA-approved ER stress and MAP kinase inhibitors *in vitro*"

**Figure S1**

**A**

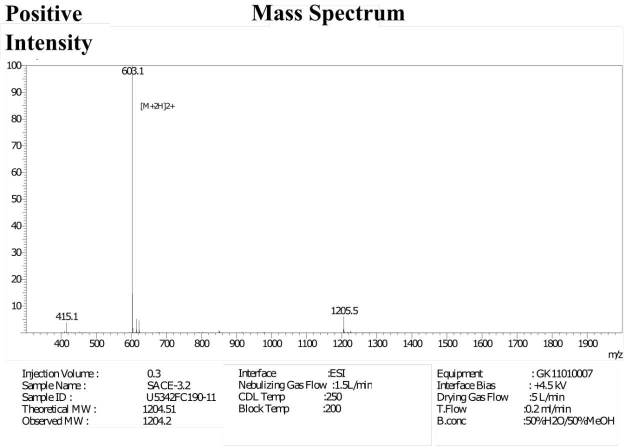


**B**

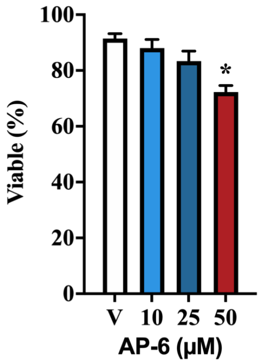


**C**

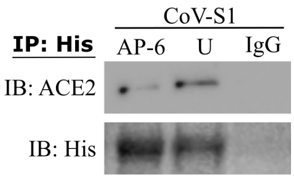


**C**

```
CoV2-S1   KIADYNYKLPDDFTGCVIAWNSNNLDSKVGGNYNYLYRLFRKSNLKPFERDISTEIYQAG
 CoV-S1   VIADYNYKLPDDFMGCVLAWNTRNIDATSTGNYNYKYRYLRHGKLRPFERDISNVPFSPD
           ************ ***:***:.*:*:.  ***** ** :*:.:*:*******.  :. .

                            AP-6 contact region
CoV2-S1   STPCNGVEGFNCYFPLQSYGFQPTNGVGYQPYRVVVLSFELLHAPATVCGPKKSTNLVKN
 CoV-S1   GKPCTP-PALNCYWPLNDYGFYTTTGIGYQPYRVVVLSFELLNAPATVCGPKLSTDLIKN
          ..**.   .:***:**:.***  *.*:***************:********* **:*:**

CoV2-S1   KCVNFNFNGLTGTGVLTESNKKFLPFQQFGRDIADTTDAVRDPQTLEILDITPCSFGGVS
 CoV-S1   QCVNFNFNGLTGTGVLTPSSKRFQPFQQFGRDVSDFTDSVRDPKTSEILDISPCSFGGVS
          :**************** *.*:* ********::* **:****:* *****:********
```

**Figure. S2**

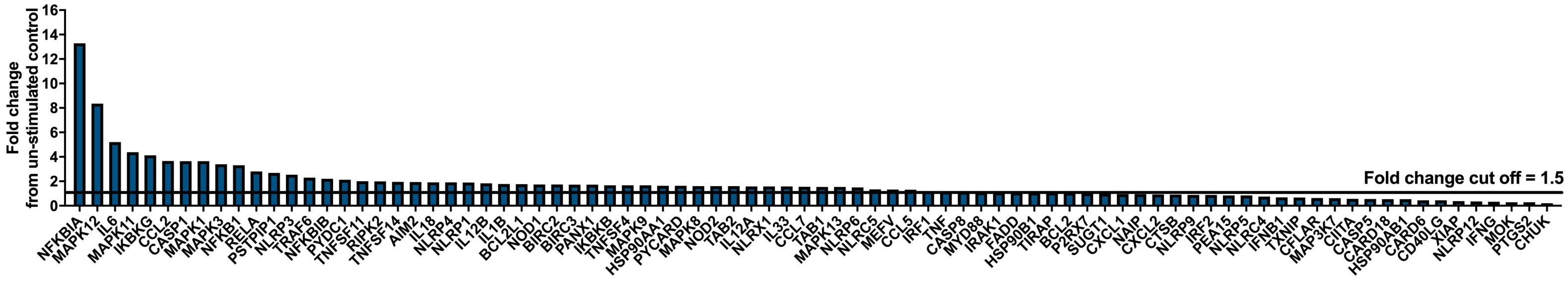
Fold change
from un-stimulated control
0
2
4
6
8
10
12
14
16
Fold change cut off = 1.5
NFKBIA
MAPK12
IL6
MAPK11
IKBKG
CCL2
CASP1
MAPK1
MAPK3
NFKB1
RELA
PSTPIP1
NLRP3
TRAF6
NFKBIB
PYDC1
TNFSF11
RIPK2
TNFSF14
AIM2
IL18
NLRP4
NLRP1
IL12B
IL1B
BCL2L1
NOD1
BIRC2
BIRC3
PANX1
IKBKB
TNFSF4
MAPK9
HSP90AA1
PYCARD
MAPK8
NOD2
TAB2
IL12A
NLRX1
IL33
CCL7
TAB1
MAPK13
NLRP6
NLRC5
MEFV
CCL5
IRF1
TNF
CASP8
MYD88
IRAK1
FADD
HSP90B1
TIRAP
BCL2
P2RX7
SUGT1
CXCL1
NAIP
CXCL2
CTSB
NLRP9
IRF2
PEA15
NLRP5
NLRC4
IFNB1
TXNIP
CFLAR
MAP3K7
CIITA
CASP5
CARD18
HSP90AB1
CARD6
CD40LG
XIAP
NLRP12
IFNG
MOK
PTGS2
CHUK

**Figure S3**

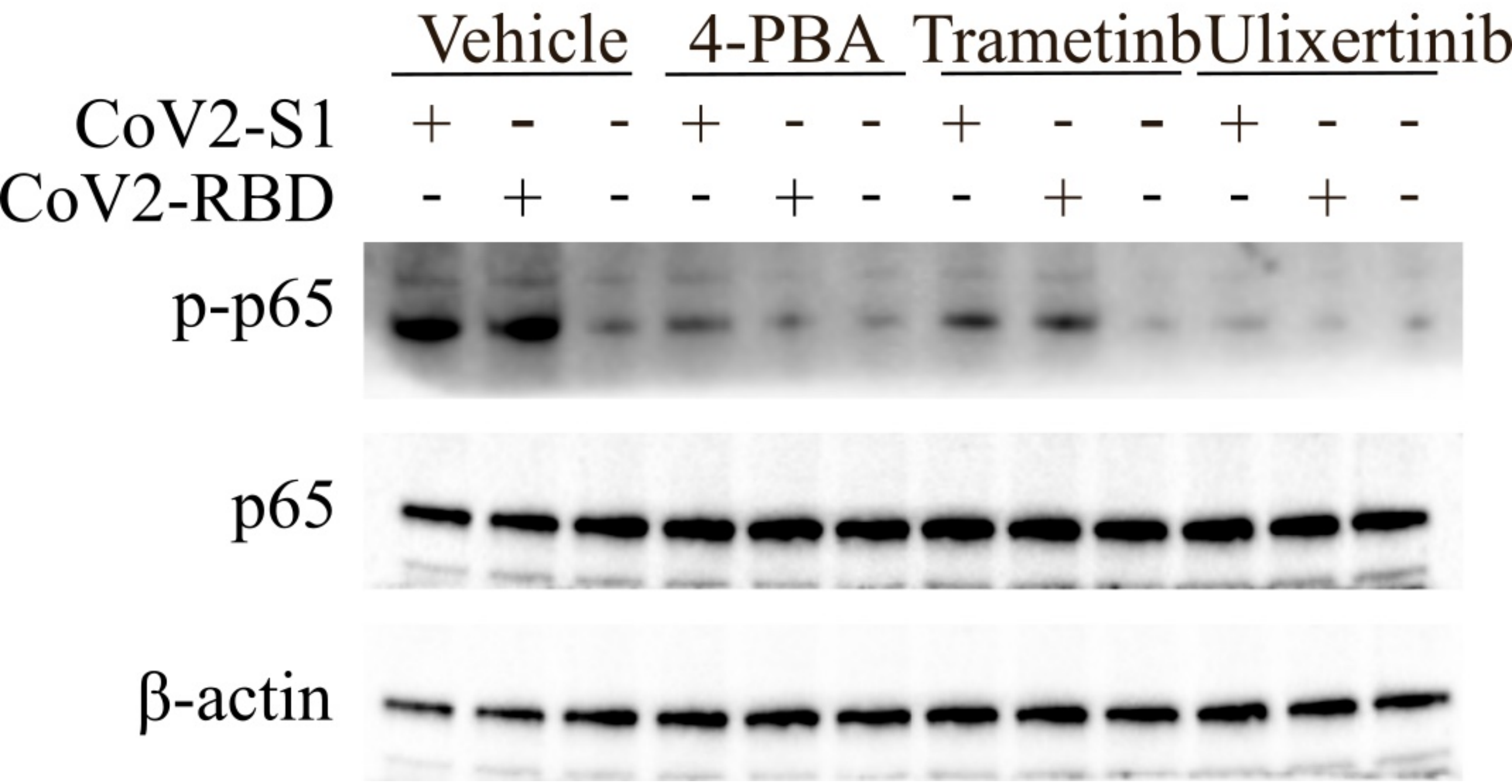
Vehicle
4-PBA
Trametinb
Ulixertinib
CoV2-S1 + - - + - - + - - + - -
CoV2-RBD - + - - + - - + - - + -
p-p65
p65
β-actin

Figure S4

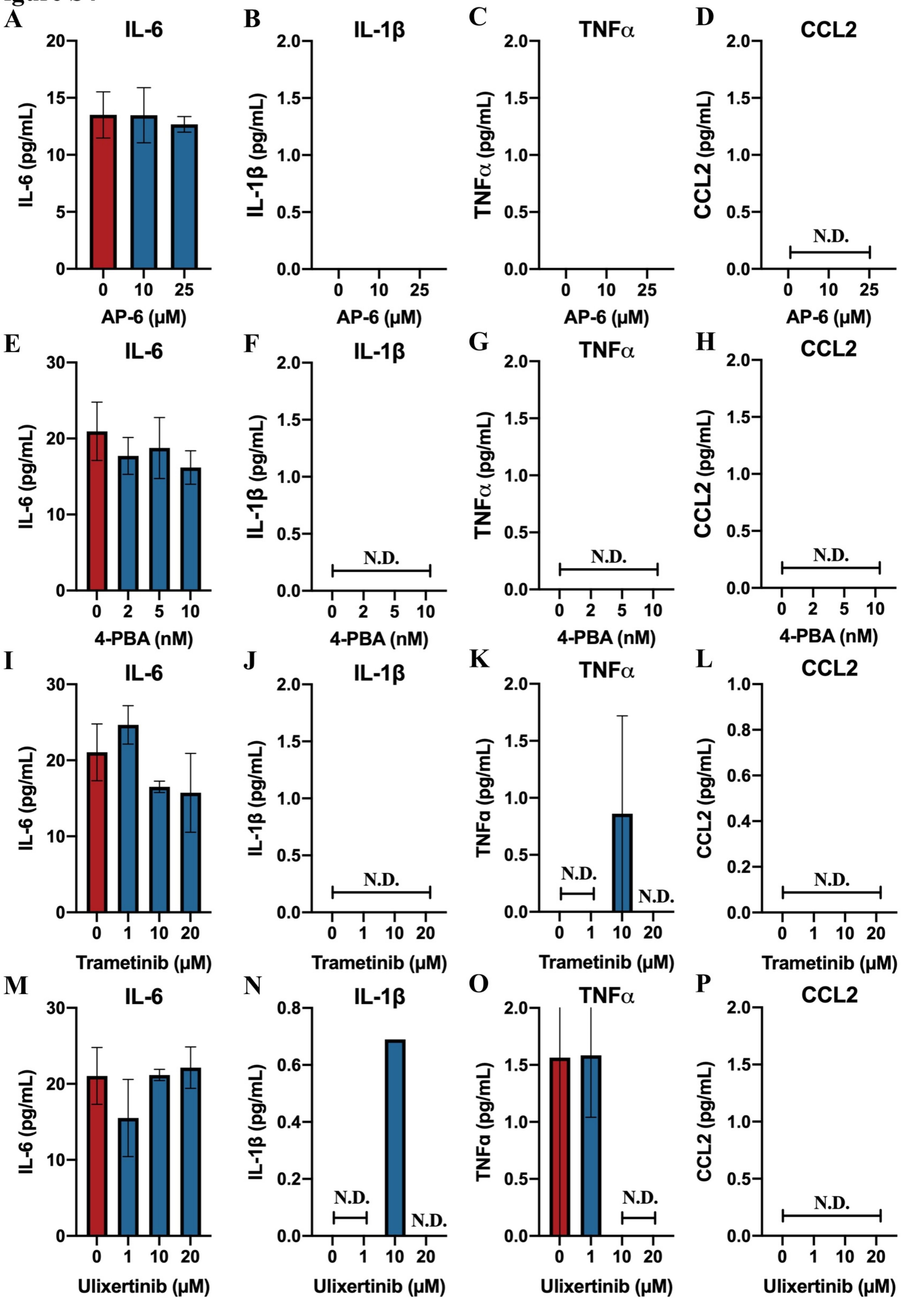

A
IL-6
IL-6 (pg/mL)
AP-6 (μM)
B
IL-1β
IL-1β (pg/mL)
AP-6 (μM)
C
TNFα
TNFα (pg/mL)
AP-6 (μM)
D
CCL2
CCL2 (pg/mL)
N.D.
AP-6 (μM)
E
IL-6
IL-6 (pg/mL)
4-PBA (nM)
F
IL-1β
IL-1β (pg/mL)
N.D.
4-PBA (nM)
G
TNFα
TNFα (pg/mL)
N.D.
4-PBA (nM)
H
CCL2
CCL2 (pg/mL)
N.D.
4-PBA (nM)
I
IL-6
IL-6 (pg/mL)
Trametinib (μM)
J
IL-1β
IL-1β (pg/mL)
N.D.
Trametinib (μM)
K
TNFα
TNFα (pg/mL)
N.D.
N.D.
Trametinib (μM)
L
CCL2
CCL2 (pg/mL)
N.D.
Trametinib (μM)
M
IL-6
IL-6 (pg/mL)
Ulixertinib (μM)
N
IL-1β
IL-1β (pg/mL)
N.D.
N.D.
Ulixertinib (μM)
O
TNFα
TNFα (pg/mL)
N.D.
Ulixertinib (μM)
P
CCL2
CCL2 (pg/mL)
N.D.
Ulixertinib (μM)

**Figure S5**

**A**

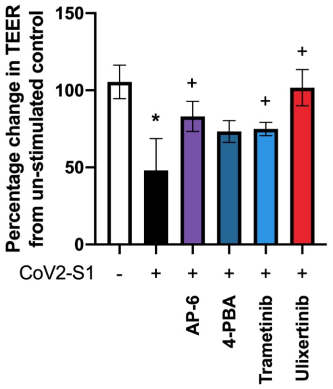

Percentage change in TEER from un-stimulated control
150
100
50
0
*
+
+
+
CoV2-S1
-
+
+
+
+
+
AP-6
4-PBA
Trametinib
Ulixertinib


**B**

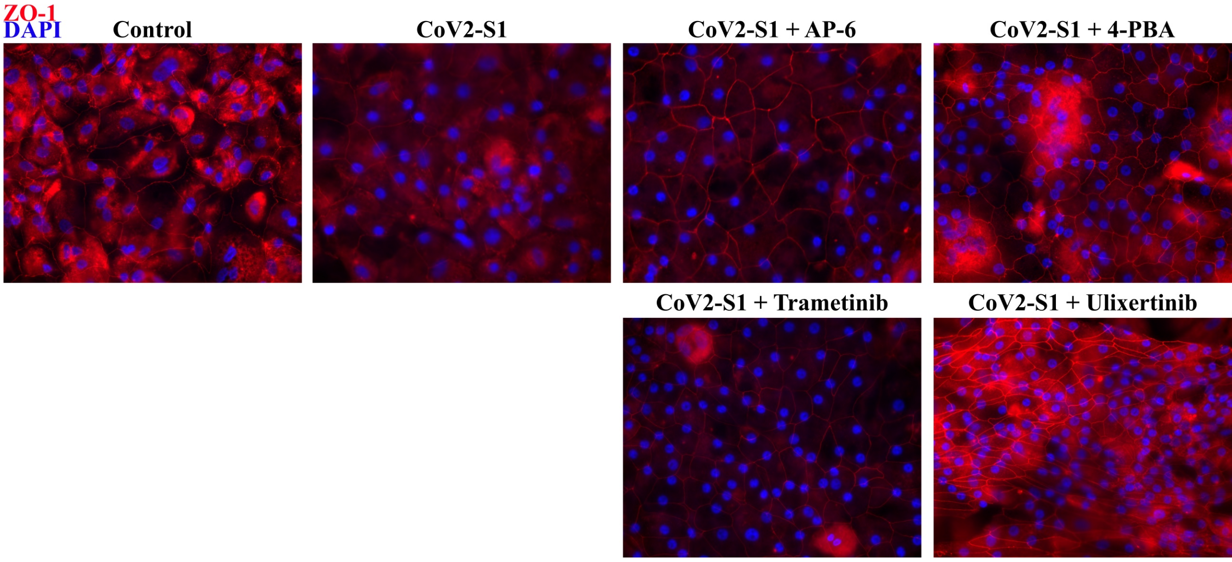

ZO-1
DAPI
Control
CoV2-S1
CoV2-S1 + AP-6
CoV2-S1 + 4-PBA
CoV2-S1 + Trametinib
CoV2-S1 + Ulixertinib
